# BET protein inhibition regulates macrophage chromatin accessibility and microbiota-dependent colitis

**DOI:** 10.1101/2021.07.15.452570

**Authors:** Michelle Hoffner O’Connor, Ana Berglind, Meaghan M. Kennedy, Benjamin P. Keith, Zachary J. Lynch, Matthew R. Schaner, Erin C. Steinbach, Jeremy Herzog, Omar K. Trad, William R. Jeck, Janelle C. Arthur, R. Balfour Sartor, Terrence S. Furey, Shehzad Z. Sheikh

## Abstract

**Introduction:** In colitis, macrophage functionality is altered compared to homeostatic conditions. Loss of IL-10 signaling results in an inappropriate and chronic inflammatory response to bacterial stimulation. It remains unknown if inhibition of bromodomain and extra-terminal domain (BET) proteins alters usage of DNA regulatory elements responsible for driving inflammatory gene expression. We determined if the BET inhibitor, (+)-JQ1, could suppress inflammatory activation of macrophages in *Il10*^-/-^ mice.

**Methods:** We performed ATAC-seq and RNA-seq on *Il10*^-/-^ bone marrow-derived macrophages (BMDMs) cultured in the presence or absence of lipopolysaccharide (LPS) and with or without treatment with (+)-JQ1 and evaluated changes in chromatin accessibility and gene expression. Germ-free *Il10*^-/-^ mice were treated with (+)-JQ1, colonized with fecal slurries and underwent histological and molecular evaluation 14-days post colonization.

**Results:** Treatment with (+)-JQ1 suppressed LPS-induced changes in chromatin at distal regulatory elements associated with inflammatory genes, particularly in regions that contain motifs for AP-1 and IRF transcription factors. This resulted in the attenuation of inflammatory gene expression. Treatment with (+)-JQ1 *in vivo* reduced severity of colitis as compared with vehicle-treated mice.

**Conclusion:** We identified the mechanism of action associated with a new class of compounds that may mitigate aberrant macrophage responses to bacteria in colitis.

**Funding:** This work was funded in part through Helmsley Charitable Trust (SHARE Project 2), NIDDK P01DK094779, NIDDK 1R01DK104828, NIDDK P30-DK034987, NIDDK 1R01DK124617, NIH Ruth L. Kirschstein National Research Service Award Individual Predoctoral Fellowship (1F31DK122704), NIH T32 Genetics NIGMS Training Grant (T32-GM007092-43), NIH T32 Translational Medicine Training Grant (T32-GM122741), NIH T32 Gastroenterology Research Training Grant (T32-DK007737), and Crohn’s and Colitis Foundation Student Research Fellowship Award, and Gnotobiotic Animal Facility.

## Introduction

Macrophages are key innate immune cells that eliminate bacteria, foreign antigens, and debris through phagocytosis and play a central role in mediating adaptive immune responses (1, 2). Macrophages maintain homeostasis in various tissues including the brain, lungs, liver, spleen, peritoneal cavity, and intestines, but these roles can vary (3). Signals from the local environment determine tissue-specific macrophage function under homeostasis (4, 5). For example, macrophages found within the intestine are tolerized to the enteric microbiota, robustly phagocytosing bacteria without releasing potent inflammatory cytokines or mediating T_H_1 or T_H_17 responses (6, 7). Changes in the local environment or presence of stimuli prompt rapid activation of macrophages, in which they gain additional functions required to help restore homeostasis (4, 5, 8).

DNA regulatory elements (DREs), including enhancers, promoters, and insulators, significantly contribute to the gene expression program in and function of macrophages (4, 5, 8, 9). Using Chromatin Immuno-Precipitation (ChIP)-seq and Assay for Transposase-Accessible Chromatin (ATAC)-seq to examine the distribution of H3K27ac modifications and nucleosome-depleted accessible chromatin, respectively, Lavin, *et al*, determined that tissue resident macrophages have unique enhancer landscapes, resulting in the preference for a distinct set of transcription factors in a tissue-dependent manner (4). Others have demonstrated that the presence of new stimuli resulted in rapid chromatin remodeling to promote expression of relevant response genes (8, 10, 11). Stimulation with lipopolysaccharide (LPS) leads to both phosphorylation of H3S28 and increased acetylation of H3K27, promoting rapid transcription of inflammatory genes, such as *Il12β*, in bone marrow-derived macrophages (BMDMs) (10, 11). Ostuni, *et al*, determined that stimulation-dependent epigenomic memory in BMDMs was established through the formation and usage of latent enhancers, identified by *de novo* H3K4me1 and H3K27ac marks, following stimulation (8). Establishment of latent enhancers provides for a rapid and specific response upon restimulation of macrophages (8). Together these studies demonstrated that the macrophage epigenetic state, under homeostasis and in response to stimuli, plays a central role in dictating macrophage function. However, epigenetic disturbances resulting in dysregulated macrophage function are understudied.

The inflammatory bowel diseases (IBDs), Crohn’s disease (CD) and ulcerative colitis (UC), are chronic inflammatory diseases of the intestines driven by exaggerated immune responses toward enteric bacteria in a genetically susceptible host (12). Macrophage loss of tolerance toward bacteria is a key event attributed to the pathogenesis of CD (13–24). We previously identified altered chromatin remodeling in macrophages from both colitis-prone *Il10*^-/-^ mice and colonic tissue isolated from CD patients (9). In wild-type (WT) and *Il10*^-/-^ BMDMs in the presence and absence of LPS stimulation as well as lamina propria macrophages (LP MΦs) from the colon of WT and *Il10*^-/-^ mice raised under germ free (GF) or specific pathogen free (SPF) conditions revealed bacteria-dependent and genotype-specific changes in chromatin accessibility. Increased chromatin accessibility was most frequently associated with LPS stimulation in BMDMs, while increased chromatin accessibility in LP MΦs was most frequently more associated with the loss of IL-10. Interestingly, the accessible chromatin regions associated with these two categories corresponded to a common set of inflammatory genes, including *Nos2, Irf1, Il7r*, and *Rel*. These data emphasize genetic predisposition as both a bacteria-independent and -dependent determinant of chromatin structure that is independent of bacteria in addition to unique chromatin responses driven by bacterial stimuli. A major challenge remains in how to prevent the formation of an aberrant chromatin state that promotes chronic intestinal inflammation in macrophages.

The bromodomain and extra-terminal domain (BET) protein family consists of four distinct proteins that recognize and also directly bind acetylated lysine residues on histone proteins, thereby contributing to gene regulatory activity (25). Small molecule binding of the BET proteins’ two bromodomain active sites prevents acetylated lysine recognition, resulting in attenuated inflammatory responses (26, 27). Nicodeme *et al* originally demonstrated BET protein binding increased upon LPS stimulation of BMDMs, but the addition of the small molecule, I-BET151, was sufficient to decrease BET protein binding and decrease expression of LPS responsive genes (26). Studies using other BET inhibitors, such as (+)-JQ1, confirmed that inhibition of BET proteins attenuates LPS- and IFN-y-induced secretion of inflammatory cytokines in BMDMs and *in vivo* murine models of endotoxic shock (27–30). Although BET proteins have been implicated in direct remodeling of chromatin, previous studies have not examined the full of extent of their role shaping chromatin accessibility in the LPS response (31, 32). Additionally, Cheung, *et al*, suggested that BET inhibitors ameliorate adoptive T-cell transfer-induced colitis, by limiting T_H_1 and T_H_17 cell differentiation (33). However, it is unclear if macrophages play a role in this process, as the mechanism by which BET inhibitors function to limit inflammatory gene expression is not entirely clear.

In this study, we sought to determine if inhibition of BET proteins using (+)-JQ1 mitigates chromatin remodeling associated with the LPS-induced inflammatory state in macrophages. We report that treatment of macrophages with (+)-JQ1 prior to LPS stimulation limited genome-wide changes in chromatin accessibility, particularly at regions distal to transcription start sites. Concomitant changes in gene expression were also observed, where 10 distinct gene expression classes were observed based on the effects of both (+)-JQ1 treatment and LPS stimulation. (+)- JQ1 attenuated the expression of approximately 90% of LPS-induced genes in macrophages. Chromatin regions distal to transcription start sites (TSSs) of LPS-induced genes revealed enrichment for AP-1 and IRF transcription factor binding motifs. (+)-JQ1 treatment prior to LPS stimulation of macrophages prevented chromatin remodeling at approximately 1,100 AP-1 and IRF motif target sequences. To evaluate efficacy of (+)-JQ1 *in vivo*, we treated GF *Il10*^-/-^ colonized with normal enteric microbiota with (+)-JQ1 and determined that (+)-JQ1 mitigates severity of colitis. Together, these data highlight that BET protein inhibition by (+)-JQ1 reduced LPS-induced chromatin remodeling in macrophages, giving rise to attenuated inflammatory responses in *Il10*^-/-^ macrophages and colitis development *in vivo*.

## Results

### Inhibition of BET proteins limits LPS-induced changes in macrophage chromatin accessibility genome-wide

Genome-wide epigenetic changes, including post-translational histone modifications and chromatin accessibility, upon LPS stimulation in macrophages have been heavily annotated (8–10, 34). Increased BET protein binding is associated with expression of LPS-induced inflammatory genes (26–29, 33). To comprehensively determine the role of BET proteins in LPS-induced chromatin remodeling, we treated *Il10*^-/-^ BMDMs with (+)-JQ1 for 12 hours prior to stimulation with LPS and evaluated changes in chromatin accessibility by ATAC-seq 4 hours later (Figure 1A). Treatment with (+)-JQ1 did not affect cell viability (Supplementary Figure 1A) or cell cycle progression (Supplementary Figure 1B).

**Figure 1.**
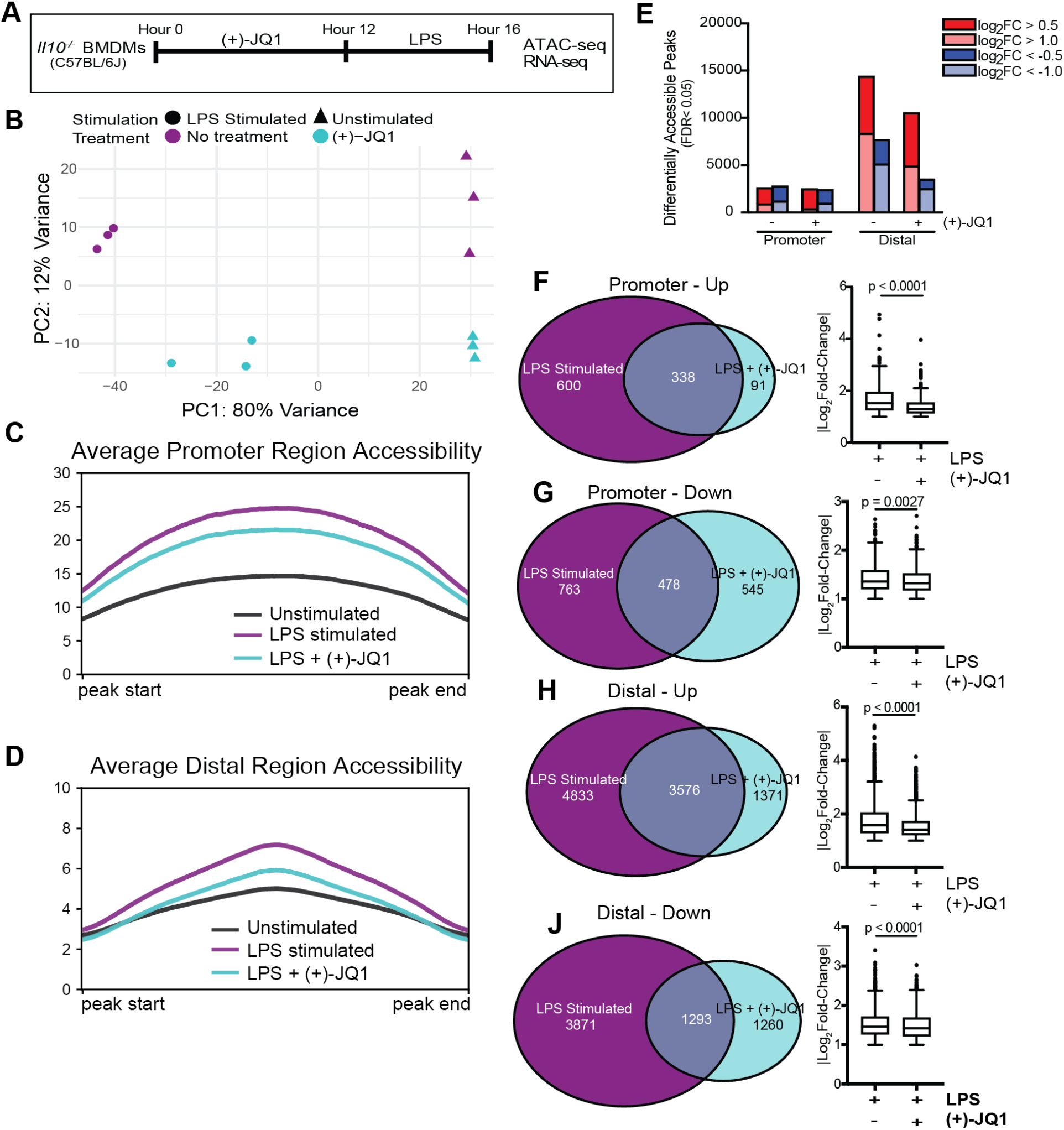
(+)-JQ1 treatment prevents LPS-induced chromatin accessibility changes at distal regions. (A) Schematic of experimental workflow. (B) Principal component analysis (PCA) of chromatin accessibility profiles for samples that remained unstimulated (triangles) or were stimulated with LPS (circles) in the absence (purple) or presence (aqua) of (+)-JQ1. Average chromatin accessibility signal profiles for (C) promoter regions (−1000/+100 bases) and (D) distal regions for unstimulated (black), LPS stimulated (purple) and LPS + (+)-JQ1 (aqua) samples. (E) Identification of regions that differentially increase (red) or decrease (blue) in accessibility during LPS stimulation with or without (+)-JQ1 as compared with unstimulated BMDMs at promoter (left) and distal (right) regions (FDR < 0.05). Identification of peaks shared with |log_2_ fold-change >1| (left) and comparison of average log_2_ fold-change values using Wilcoxon sum rank tests for (F) promoters increasing in accessibility, (G) promoters decreasing in accessibility, (H) distal regions increasing in, and (J) distal regions decreasing in accessibility. FDR adjusted P values determined using DESeq2. N=3 biological replicates.

Principle component analysis (PCA) of chromatin accessibility profiles in *Il10*^-/-^ BMDMs (Figure 1B) showed the first principal component (PC1) stratified samples based on LPS stimulation status, demonstrating that LPS was the primary driver of chromatin accessibility changes. Samples were further segregated along PC2 based on treatment with (+)-JQ1. Accessibility profiles from samples treated with (+)-JQ1 prior to LPS stimulation were more similar to those from unstimulated samples along PC1 than compared to untreated, LPS stimulated samples (Figure 1B), suggesting (+)-JQ1 mitigated some LPS-induced chromatin accessibility changes in *Il10*^-/-^ BMDMs.

Both LPS stimulation and the loss of IL-10 expression were previously shown to affect genomewide chromatin accessibility in *Il10*^-/-^ BMDMs (9). To determine how (+)-JQ1 modifies genomewide LPS-stimulated chromatin accessibility, we determined the average chromatin accessibility at both promoter (n = 36,670, Figure 1C) and distal (n=157,614, Figure 1D) sites and quantified the number and magnitude of differentially accessible regions (DARs; FDR < 0.05) in LPS-stimulated and LPS + (+)-JQ1 conditions, compared to unstimulated, *Il10*^-/-^ BMDMs (Figure 1E). Average chromatin accessibility at both promoter and distal sites was decreased with (+)-JQ1 treatment compared to LPS stimulation alone (Figures 1C & 1D). However, the relative decrease in accessibility was greater at distal sites compared to promoter sites. Approximately 15% of the total number of promoter and distal sites were differentially accessible (Figure 1E). The overall number of DARs (both increased and decreased) at promoter sites were roughly equivalent with or without (+)-JQ1 treatment, but a greater percentage of DARs without (+)-JQ1 treatment were high magnitude changes (|log_2_ fold-change(FC)| > 1; Figure 1E). In contrast, at distal sites, both the overall number of DARs as well as the percentage of high magnitude DARs were decreased substantially with (+)-JQ1 (Figure 1E). DARs with high magnitude changes at both promoter and distal sites demonstrated greater concordance between the LPS and LPS + (+)-JQ1 conditions when accessibility was increased by LPS stimulation than when decreased (Figures 1F-1J). Even amongst these DARs, (+)-JQ1 treatment significantly decreased the average magnitude (|log_2_FC|) for all four categories (Figures 1F-1J, right; Wilcoxon sum rank test, p < 0.005). Based on these data, we conclude that (+)-JQ1 treatment attenuates the effect of LPS on genome-wide chromatin accessibility and has a greater impact on DARs at distal sites.

### Exogenous IL-10 treatment drives changes in chromatin accessibility distinct from BET inhibition

We previously showed that supplementation with IL-10 had minimal effects on genome-wide chromatin remodeling in *Il10*^-/-^ BMDMs (9). To determine how IL-10 treatment compared to (+)- JQ1 in limiting LPS-induced chromatin remodeling, we treated *Il10*^-/-^ BMDMs with IL-10 for 12 hours followed by LPS stimulation for 4 hours and ATAC-seq. PCA revealed that chromatin accessibility profiles from samples treated with IL-10 clustered separately from samples treated with (+)-JQ1 and untreated samples, regardless of LPS stimulation status (Supplementary Figure 2A). IL-10 treated *Il10*^-/-^ BMDMs showed decreased promoter accessibility compared with (+)-JQ1 treated samples, on average (Supplementary Figure 2B), while (+)-JQ1 treated samples had decreased accessibility at distal regions compared to IL-10 treated samples (Supplementary Figure 2C). Compared to unstimulated and untreated *Il10*^-/-^ BMDMs, IL-10 treatment prior to LPS stimulation resulted in less promoter DARs than either LPS stimulation alone or with (+)-JQ1 treatment (Supplementary Figure 2D). In contrast, the number of DARs (increased and decreased) detected at distal sites with IL-10 treatment was similar to what was observed with LPS stimulation alone (Supplementary Figure 2D). To quantify the effects of IL-10 or (+)-JQ1 treatment on LPS stimulated chromatin accessibility, we correlated normalized signals across all accessible regions between LPS stimulated alone, LPS + (+)-JQ1, and LPS + IL-10 and unstimulated *Il10*^-/-^ BMDMs (Supplementary Figures 2E-G). We found (+)-JQ1 treated samples correlated best with unstimulated samples, suggesting a greater impact of (+)-JQ1 than IL-10 on limiting LPS-induced chromatin remodeling. Together, these data show (+)-JQ1 better preserves a non-inflammatory chromatin state in LPS-stimulated *Il10*^-/-^ BMDMs, and IL-10 signaling affects LPS-induced chromatin remodeling distinctly from (+)-JQ1.

### BET protein inhibition attenuates LPS-induced expression of inflammatory genes

LPS stimulation of *Il10*^-/-^ BMDMs results in robust expression of key inflammatory genes, such as *Il12β*, than is observed in LPS stimulated WT BMDMs (20, 22). To better understand how BET protein inhibition affects LPS-induced gene expression, we performed RNA-seq in BMDMs from the same mice as above (Figure 1A). Similar to our ATAC-seq analyses, PCA on these data revealed the primary source of variation was due to LPS stimulation (PC1; Figure 2A), while (+)-JQ1 treatment contributes secondarily to variation (PC2). Again, BMDMs treated with (+)-JQ1 prior to LPS stimulation clustered closer to unstimulated BMDMs along PC1 than untreated LPS stimulated BMDMs.

**Figure 2.**
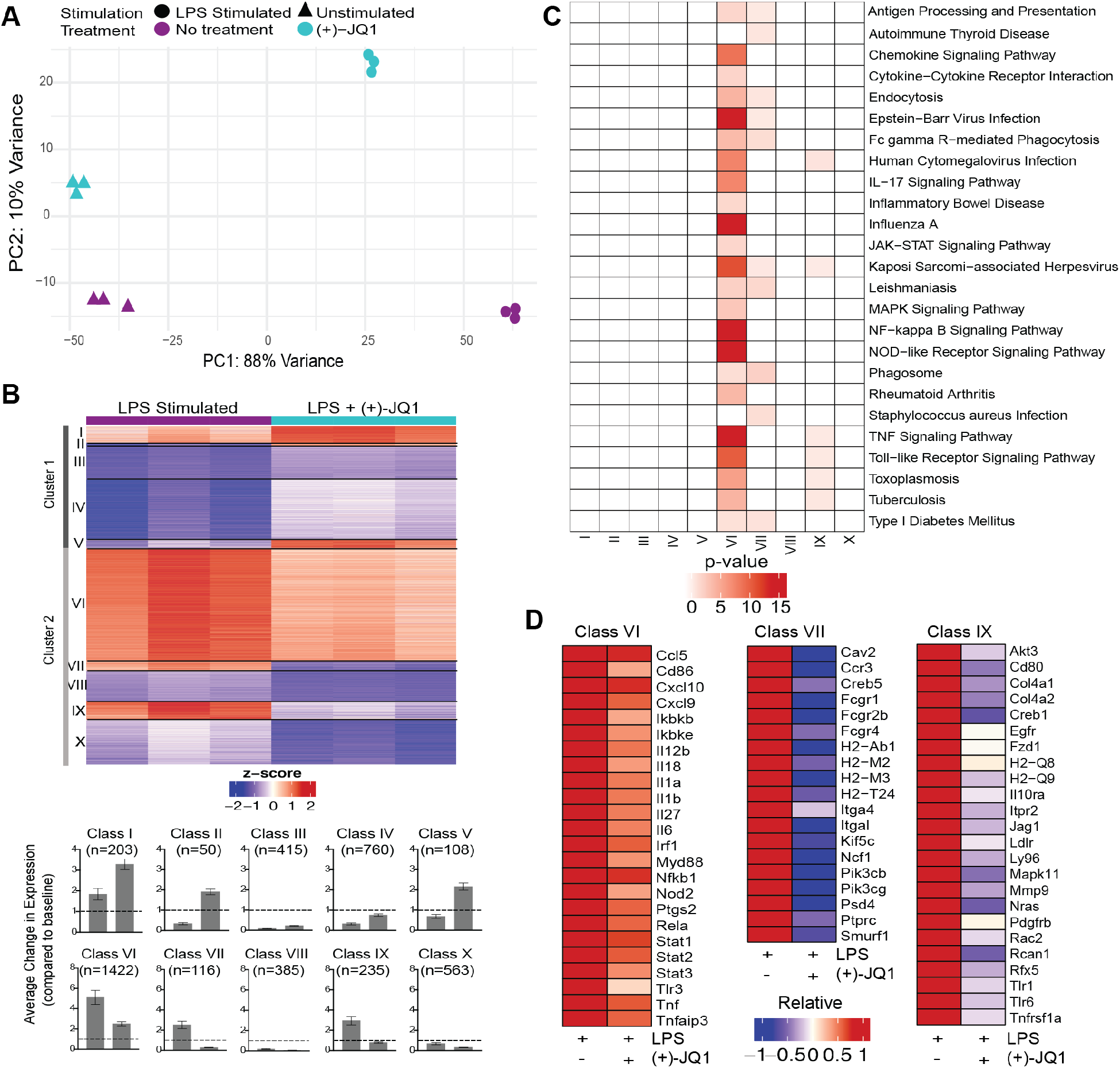
(+)-JQ1 treatment attenuates LPS-induced inflammatory gene expression in *Il10*^-/-^ macrophages. (A) PCA of gene expression profiles for samples that remained unstimulated (triangles) or were stimulated with LPS (circles) in the absence (purple) or presence (aqua) of (+)- JQ1. (B) Representative heatmap of calculated z-scores based on log_2_ fold-change of individual replicates for the LPS and LPS + (+)-JQ1 conditions normalized to the average of unstimulated, untreated *Il10*^-/-^ BMDMs. Genes were identified and divided into 2 major clusters and sub-divided into 5 classes through differential analyses outlined in Supplementary Table 1. Bar graphs represent average change in expression for LPS and LPS (+)-JQ1 conditions normalized to baseline. (C) KEGG pathway analysis for the genes identified in the 10 classes in (B). (D) Heatmaps of representative genes from Classes VI, VII, and IX that correspond to genes induced by LPS that are attenuated with (+)-JQ1. N=3 biological replicates.

To determine which genes were affected by LPS stimulation both with and without (+)-JQ1 treatment, we performed pairwise comparisons of each experimental condition with unstimulated samples. We identified 10,656 differentially expressed genes (DEGs) present in at least one of these comparisons. We then used a likelihood ratio test (LRT) to determine which genes were significantly affected by (+)-JQ1 during LPS stimulation, resulting in identification of 4,257 DEGs to further analyze.

These DEGs were clustered based on the direction of effect of (+)-JQ1 treatment compared with no treatment. Cluster 1 (n = 1,536 genes) contained genes with higher expression after LPS stimulation when pretreated with (+)-JQ1, while Cluster 2 (n = 2,721 genes) contained those with lower relative expression (Figure 2B). Within each cluster we identified five distinct classes based on adjusted p-value and magnitude of log_2_FC from comparison with unstimulated samples (Supplementary Table 1). In Cluster 1, three small classes contained genes that were significantly up-regulated by LPS with (+)-JQ1 treatment (log_2_FC > 1) but were either weakly induced by (Class I; n = 203 genes), down-regulated by (Class II; n = 50 genes), or showed minimal response to (Class V; n = 108 genes) LPS stimulation alone. In the other two larger classes (Class III, n = 415 gene; Class IV, n = 760 genes), (+)-JQ1 treatment resulted in relatively higher expression compared to LPS stimulation alone, but expression levels of these genes were all below those in unstimulated BMDMs (Figure 2B). Pathway enrichment analysis of Cluster 1 genes revealed enrichment for Longevity Regulating, Mismatch Repair, and Systemic Lupus Erythematosus pathways, but no immune related pathways (Supplementary Figure 3A).

In Cluster 2, which consists of genes whose relative expression is lower when treated with (+)- JQ1 prior to LPS stimulation (Figure 2B), expression of nearly all (~90%) of the genes was significantly induced during LPS stimulation alone (log_2_FC > 1). The most robustly up-regulated genes formed Class VI (n = 1,422 genes). Class VI contained genes upregulated by LPS stimulation regardless of (+)-JQ1 treatment but where pre-treating with (+)-JQ1 resulted in an average 2-fold decrease in expression compared with LPS stimulation alone. Pathway analysis using these genes showed enrichment for more than 20 pathways associated with inflammatory and bacterial response (Figure 2C) and included many pro-inflammatory genes such as *Il12β, Il6*, and *Tnf* (Figure 2D). Class VII (n = 116 genes) and Class IX (n = 235 genes) consisted of genes induced by LPS stimulation alone, but whose expression levels when treated with (+)-JQ1 treatment were lower than or similar to, respectively, expression in unstimulated *Il10*^-/-^ BMDMs (Figure 2B). Class VII genes were enriched for macrophage functions, such as antigen presentation and phagocytosis, while Class IX genes were involved in bacterial and viral response pathways and TNF signaling (Figure 2C). The remaining two classes (Class VIII, n = 385 genes; Class X, n = 563 genes) contained genes that were down-regulated upon LPS stimulation regardless of (+)-JQ1 treatment, but whose expression was reduced to a greater extent when pretreated with (+)-JQ1 (Figure 2B). Neither class showed enrichment for immune pathways (Figure 2C).

### Exogenous IL-10 exerts limited anti-inflammatory effects on LPS-induced gene expression

IL-10 negatively regulates expression of *Il12β* and other inflammatory genes in macrophages (20, 22). We wanted to better understand how exogenous IL-10 affects the *Il10*^-/-^ BMDM response to LPS and to compare this with (+)-JQ1 treatment. PCA revealed that IL-10 treatment prior to LPS stimulation resulted in transcriptional profiles markedly similar to untreated LPS stimulated BMDMs (Supplementary Figure 3B), including across genes in our ten previously defined LPS response classes (Supplementary Figure 3C). Expression fold-changes upon LPS stimulation of IL-10 treated samples varied across key inflammatory genes found in Class VI. Expression of some genes, such as *Il12β*, was attenuated compared to (+)-JQ1 treated samples; the majority of other genes, such *Nod2*, showed increased expression compared with (+)-JQ1 treatment and was more similar to untreated samples (Supplementary Figures 3D and 3E). In contrast, IL-10 treatment had little effect on LPS-induced Class VII or IX gene expression (Supplementary Figures 3D and 3E). Collectively, these results demonstrate that (+)-JQ1 treatment has a strong effect on genes induced by LPS stimulation, which were more extensive than with IL-10 treatment and resulted in down-regulation of genes implicated in the inflammatory response.

### (+)-JQ1 restricts chromatin accessibility at distal regulatory elements of Class VI genes

Distal regulatory elements, such as enhancers, largely contribute to gene expression programs that define cell identity (4, 5). Recent studies have suggested that BET protein inhibition alters use of cell-specific super-enhancers (34–36). Therefore, we sought to determine the effects of (+)-JQ1 on chromatin accessibility and altered expression at LPS-responsive genes. Across our ten gene expression classes, we found a similar number of distal accessible regions near member genes on average (+/- 25kb from the TSS; Supplementary Figure 4A, 4B). LPS stimulation increased distal chromatin accessibility in all gene classes, but the magnitude varied (Supplementary Figure 4C). Consistent with our genome-wide analyses, (+)-JQ1 treatment decreased the effect of LPS stimulation on chromatin accessibility changes for genes of all 10 classes but most notably for genes in Classes VI, VII, and IX that contain genes with attenuated expression (Figures 3A & 3B; Supplementary Figure 4C). In contrast, a similar analysis of IL-10 treated samples on Class VI, VII, and IX genes showed accessibility at promoter regions was less altered on average than with (+)-JQ1 treatment (Supplementary Figure 4D). However, accessibility at distal regions associated with these classes more closely resembled those of untreated LPS stimulated samples, especially for Class VII, compared to (+)-JQ1 treatment (Supplementary Figure 4E).

**Figure 3.**
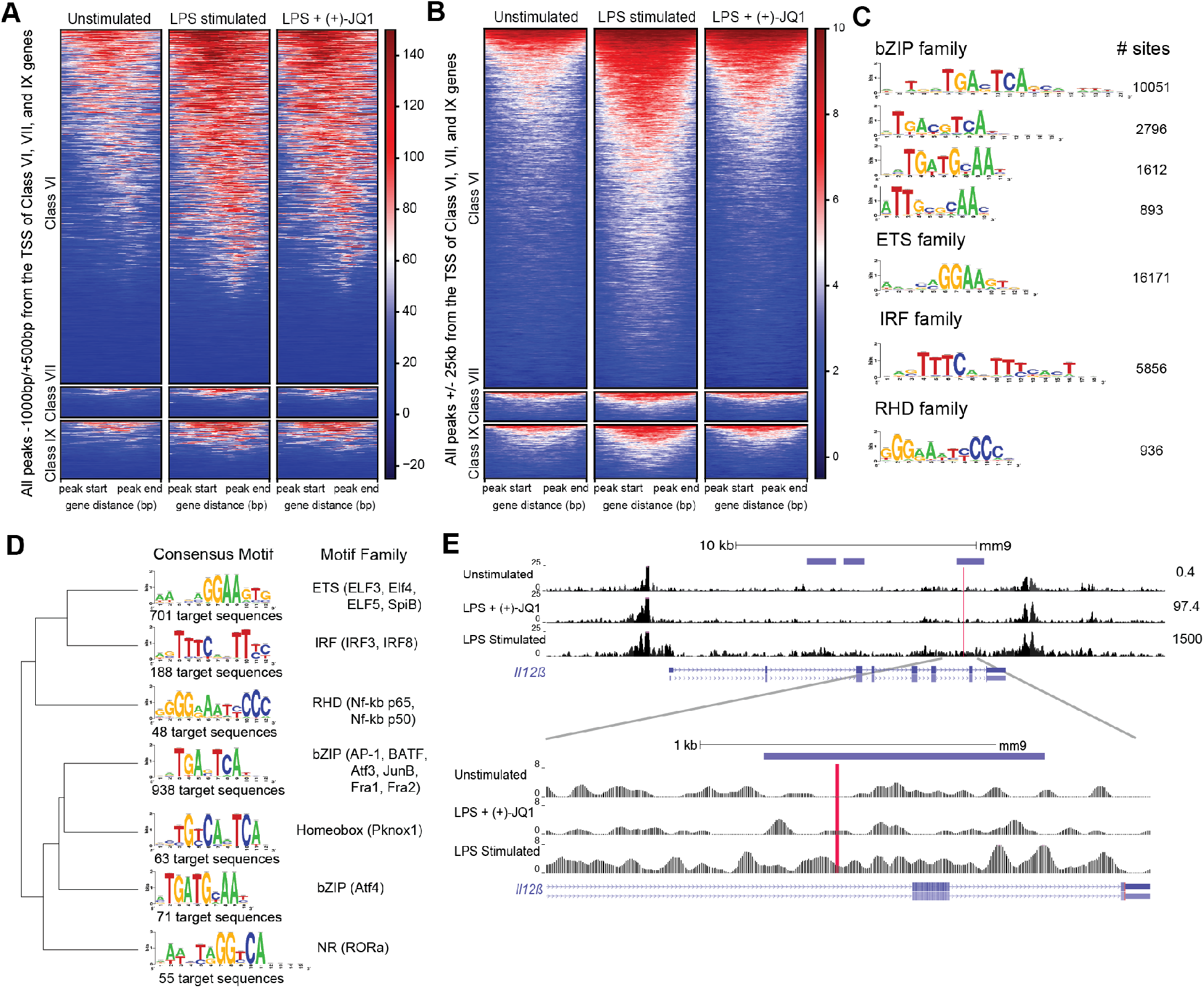
(+)-JQ1 prevents LPS-induced chromatin remodeling at putative AP-1 and IRF regulatory elements associated with class VI genes. Heatmaps of accessible chromatin from unstimulated (left), LPS stimulated (center) or LPS + (+)-JQ1 conditions for Classes VI (top), VII (center), and IX (bottom) for ChIPSeeker identified (A) promoter regions and (B) distal regions. (C) Identification and quantification of sites containing known HOMER motifs for regions that increase in accessibility with LPS stimulation compared to unstimulated, untreated BMDMs (FDR < 0.05 for differential analysis, FDR < 0.10 for motif enrichment). (D) Identification, motif clustering, and quantification of sites containing known HOMER motifs for regions that decrease in accessibility with (+)-JQ1 treatment prior to LPS stimulation compared to LPS stimulation alone (FDR < 0.05 for differential analysis, FDR < 0.10 for motif enrichment). (E) Gene tracks of ATAC-seq peaks within 25kb of the TSS of *Il12β* for unstimulated (top), LPS + (+)-JQ1 (center) and LPS stimulated (bottom) conditions. Horizontal purple bars above the tracks are representative of peaks that are differentially increased during LPS stimulation compared to unstimulated BMDMs and are differentially decreased with (+)-JQ1 compared to LPS alone (FDR < 0.05). Red line is representative of putative AP-1 motif identified by HOMER. Numbers to the right represent corresponding *Il12β* transcript levels. FDR adjusted P values for differential analysis determined using DESeq2 and for motifs using HOMER. N=3 biological replicates.

Focusing more specifically on regions that significantly changed upon LPS stimulation, we found an enrichment for DARs near genes from Classes VI (n=504) and VII (n=47), but not Class IX (n=66) (hypergeometric test; p = 9.22×10^−16^, p = 7.56×10^−4^, p = 0.24, respectively). As Class VI was the largest and contains key inflammatory genes, we investigated chromatin accessibility separately at promoter and distal regions. Both promoter (Figure 3A) and distal (Figure 3B) regions showed increased chromatin accessibility upon LPS stimulation compared with unstimulated *Il10*^-/-^ BMDMs. Consistent with our genome-wide observations, (+)-JQ1 treatment showed little effect on promoter regions (Figure 3A), but distally we found an overall attenuation of accessibility changes and a reduced number of DARs in response to LPS stimulation (Figure 3B). Motif analysis of distal DARs associated with Class VI genes with increased accessibility in untreated LPS stimulated samples (n = 5,323 regions) showed enrichment for motifs of bZIP transcription factors, including many AP-1 family sub-units, ETS transcription factors, including PU.1, and IRF family members (Figure 3C). DARs where (+)-JQ1 attenuated LPS-induced accessibility changes (n = 920 regions) were enriched for motifs for a subset of AP-1 family members and several IRF and RHD (NF-kB p65/p50) factors (Figures 3D & 3E). DARs with decreased accessibility during LPS stimulation alone or increased accessibility during (+)-JQ1 treatment with LPS were enriched for motifs from ETS family members (Supplementary Figure 4F & 4G).

Overall, these data suggest that (+)-JQ1 treatment restricts LPS-induced chromatin accessibility changes at distal regulatory elements near inflammatory genes. Interestingly, our data also suggests that exogenous IL-10 attenuates LPS-induced chromatin changes, but primarily at promoters of LPS-inducible genes.

### (+)-JQ1 treatment limits experimental colitis in germ free Il10^-/-^ mice

To determine whether (+)- JQ1 mitigates colitis *in vivo*, we treated adult *Il10*^-/-^ mice previously housed under GF conditions with (+)-JQ1 (I.P. injections) before colonization with SPF C57BL/6J fecal matter slurries (Figure 4A). Additional (+)-JQ1 treatments were administered on Days 3, 6, 9, and 12, and animals were sacrificed on Day 14 (Figure 4A). Treatment with (+)-JQ1 did not result in any significant changes in weight as compared with vehicle treated mice (Supplementary Figure 5A).

**Figure 4.**
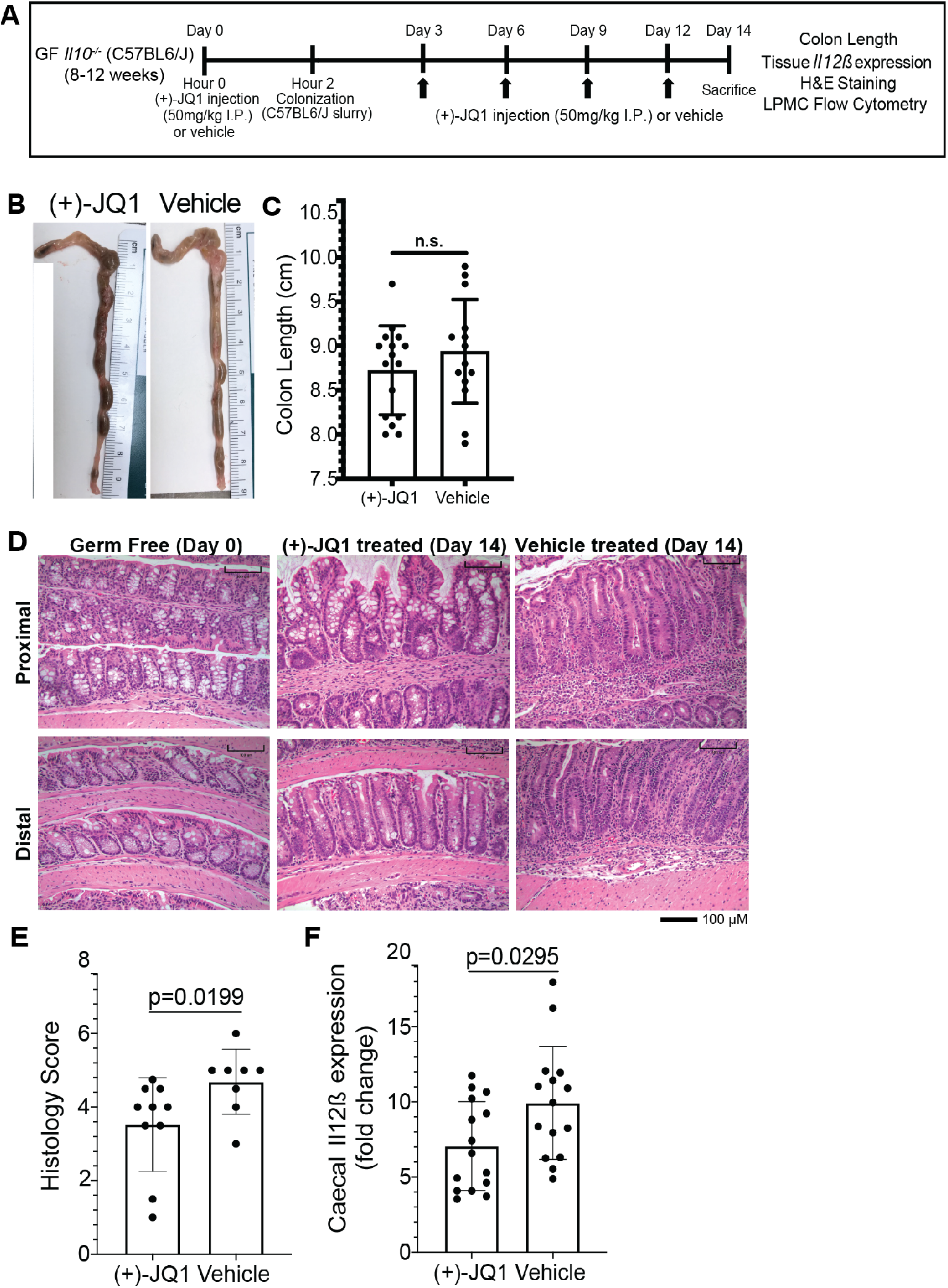
(+)-JQ1 treatment attenuates microbiota-induced colitis in germ free *II10*^-/-^ mice. (A) Schematic of experimental workflow (n=10+ mice/conditions and represent at least 3 independent experiments. (B) Representative images of colon length and thickness in (+)-JQ1- and vehicle-treated mice. (C) Quantification of average colon length (n=16 (+)-JQ1; n=15 vehicle). (D) Representative images of H&E stained colons for proximal (top) and distal (bottom) portions of the colon in germ free (GF) mice (left), colonized mice treated with (+)-JQ1 harvested on Day 14 (center) and colonized mice treated with vehicle harvested on Day 14 (right). Scale represents 100 μM. (E) Quantification of histology scores based on H&E staining (n=10 (+)-JQ1; n=8 vehicle). (F) Quantification of cecal *Il12β* expression by qPCR (n=15/treatment). Significance values determined using a 2-tailed unpaired, non-parametric student’s t test.

We isolated lamina propria mononuclear cells (LPMCs) and splenocytes from (+)-JQ1 and vehicle treated animals to identify changes in immune cell population sizes associated with treatment (Supplementary Figure 5B). Total CD45^+^ leukocytes constituted ~60% of isolated LPMCs and 75% of splenocytes in vehicle treated mice, which did not significantly change with (+)-JQ1 treatment (Supplementary Figure 5C). No significant differences in the proportion of CD11b^+^CD11c^-^F4/80^+^ macrophages or total CD3^+^ T cells and CD19^+^ B cells isolated from the intestine or spleen were observed with (+)-JQ1 treatment (Supplementary Figure 5D & 5E). Based on these observations, we conclude that the dose and frequency of (+)-JQ1 given to these mice did not have a significant effect on immune cell numbers in the colon or spleen.

Finally, we investigated the effect of (+)-JQ1 treatment on colonic inflammation in these mice. Colon length and morphology after (+)-JQ1 treatment were unchanged from vehicle treated mice (Figures 4B and 4C). Histological scoring using a modified method from Berg, *et al*, where colons from uninflamed uncolonized GF *Il10*^-/-^ mice harvested on Day 0 were used as a comparison (Figure 4D). (+)-JQ1 treated mice had significantly lower inflammation scores compared to vehicle treated mice (Figure 4E). We quantified cecal *Il12β* mRNA expression; (+)-JQ1 treatment significantly decreased relative *Il12β* expression compared to vehicle treated mice (Figure 4F). Taken together, these data demonstrate that treatment with (+)-JQ1 attenuates severity of colitis in GF *Il10*^-/-^ mice two weeks post-colonization.

## Discussion

The enhancer landscape is central to determining the gene expression program and identity of a cell (4, 5). Both the microenvironment and external stimuli influence macrophage enhancer utilization (4, 5, 8–11). Previously, we showed that putative enhancers regions are differentially accessible in a genotype- and stimulation-dependent manner in *Il10*^-/-^ macrophages (9). In this study, we report that BET protein inhibition with (+)-JQ1 is sufficient to attenuate LPS-induced chromatin remodeling at putative enhancers containing AP-1 and IRF motifs. Decreased accessibility is accompanied by attenuation of key LPS-induced inflammatory genes, whose TSSs are found in close proximity to these putative enhancers. Binding of BET proteins, particularly BRD4, to chromatin in macrophages increases following LPS stimulation (26, 34), which allows the formation of LPS-induced super-enhancers in close proximity to inflammatory genes such as *Tnf* and *Nfkbia* (34). We propose that treatment with (+)-JQ1 inhibits BET protein-mediated nucleosome eviction thus preventing LPS-induced increases in chromatin accessibility at putative enhancers driving inflammatory gene expression (32). Without ChIP-seq datasets and functional assays, we cannot confirm the regions identified in this study are enhancers. Future studies to annotate H3K4me1 and H3K27ac marks in these regions will be helpful. Additional functional analyses with luciferase-reporter and electrophoretic mobility shift assays can confirm enhancer activity and identify transcription factors associated with promoting regulatory activity at these sites.

Previous publications have focused on pairwise differential gene expression analyses in macrophages between LPS and LPS + BET inhibitor or BET knock-out conditions, and consistently highlight the relative down regulation of inflammatory genes (26, 28, 33, 34). However, gene expression profiles of unstimulated macrophages are seldom included in these studies despite the fact many inflammation-related surface receptors are expressed and central to macrophage functioning at baseline. Our study utilized an objective approach that presents a genome-wide annotation of LPS-stimulated gene expression changes and how (+)-JQ1 modulates this response relative to the expression observed in an unstimulated *Il10*^-/-^ BMDMs. While our findings regarding the attenuation of inflammatory genes with BET inhibition is consistent with prior studies, we demonstrate that the majority of genes induced by LPS stimulation still show increased expression with (+)-JQ1 treatment compared to unstimulated macrophages (Class VI). These data indicate that BET inhibition by (+)-JQ1 at levels that do not affect cell viability does not completely abrogate the ability of macrophages to respond to a bacterial stimulus. Our approach identified additional gene expression dynamics associated with (+)-JQ1 treatment, where a subset of LPS-induced genes had expression levels below that of unstimulated macrophages (Class VII) and another subset had expression levels that were comparable to unstimulated macrophages (Class IX). These genes include *Fcgr1, Fcgr2b, Fcgr4* (Class VII), *Tlr1, Tlr6*, and *Cd80* (Class IX), which are involved in antibody-mediated phagocytosis, bacterial signaling, and co-stimulation of T cells. Genomic data alone is not sufficient to determine the functions retained by macrophages in the presence of (+)-JQ1, and sparse data exist on how BET inhibition modulates macrophage functioning (37). Therefore, it is necessary for future studies to determine if macrophages retain their phagocytic and functional capabilities upon BET inhibition.

We previously proposed that IL-10 signaling may promote the early establishment of accessible regulatory elements in the developing macrophage (9). Our findings here illustrate a novel role for IL-10 in facilitating chromatin remodeling, but it remains unclear if IL-10 directly interacts with chromatin or if these changes are prompted by downstream effectors in the IL-10 pathway. Treatment of *Il10*^-/-^ BMDMs with IL-10 alters the LPS-induced chromatin landscape, which is distinct from the resulting chromatin landscape associated with BET inhibition. Chromatin profiles of samples treated with IL-10 or (+)-JQ1 prior to LPS stimulation more closely resemble unstimulated samples than untreated LPS stimulated samples, but these treatments generate distinct chromatin profiles. Interestingly, IL-10 was more potent in attenuating promoter accessibility changes, whereas (+)-JQ1 had a larger effect on distal regions. This observation, coupled with upregulation of inflammatory genes that is comparable to untreated LPS stimulated samples, suggests that IL-10 treatment is not sufficient to prevent LPS-induced chromatin changes driving upregulation of inflammatory genes. In contrast, (+)-JQ1 treatment resulted in the attenuation of LPS-induced gene expression, including genes involved in anti-inflammatory pathways promoted by IL-10, including *Stat3* and *Jak2* (38). Previously, it was shown the IL-10 serum levels were significantly lower in in an LPS-induced endotoxic shock model when mice were treated with (+)-JQ1 (27). More recent studies demonstrated that IL-10 producing B regulatory cells rely on BRD4 to promote expression of IL-10 (39). Together, these and our data suggest that BET inhibition results in the repression of key anti-inflammatory pathways, in addition to inflammatory pathways. Future experiments should further interrogate the effects of (+)-JQ1 on macrophage LPS-induced gene expression with intact IL-10 signaling to determine the potential short and long-term consequences that may result due to the unintended inhibition of a key anti-inflammatory pathway.

The ability of BET protein inhibitors to attenuate inflammation by modulating the function of macrophages in colitis is largely unexplored. Cheung, *et al*, demonstrated that treatment with BET inhibitors limited T_H_1 and T_H_17 differentiation and prevented adoptive T-cell transfer induced colitis; Weinerroither, *et al*, revealed that the use of BET inhibitors at the onset of DSS-induced colitis exacerbated the phenotype compared to DSS administration alone (33, 37). It is unclear whether the adaptive immune findings described above are due to changes we report in macrophage phenotype. The DSS model of colitis is an acute injury model mediated by intestinal epithelial cell damage suggesting that (+)-JQ1 may prevent the expansion and differentiation of intestinal stem cells, resulting in worsened tissue injury (40, 41). Our study presents the first evaluation of BET inhibitors as a method of mitigating colitis severity in a genotype-driven, microbiota-dependent mouse model. There are limitations to our study. First, while (+)-JQ1 treatment results in modest reduction in colitis severity compared to vehicle-treated mice, further studies are needed to identify the optimal dosing strategy that maximizes attenuation of inflammation without causing harm to the intestinal epithelium. Furthermore, our study focuses on prevention of inflammation onset, not as a therapeutic, necessitating additional studies that utilize (+)-JQ1 following intestinal colonization and intestinal inflammation. Finally, although we demonstrate reduced expression of *Il12β*, an innate immune cytokine known to be produced by macrophages, future genomic studies are necessary to determine whether the effect of I.P. injection of (+)-JQ1 on *Il10*^-/-^ BMDM gene expression and chromatin remodeling translates to LP MΦ *in vivo*.

To our knowledge, there are no studies that have determined if any of the chromatin-based effects of (+)-JQ1 are permanent following removal of (+)-JQ1 in culture. Gibbons, *et al*, recently demonstrated that (+)-JQ1 represses expression of IFN-γ in T_H_1 cells, but this blockade only resulted from the absence of BET protein binding at the promoter, not remodeling of chromatin associated with known enhancers for IFN-γ (30). Consequently, removal of (+)-JQ1 from culture resulted in the restoration of IFN-γ expression. In contrast, other studies have demonstrated blockade of chromatin modifiers is sufficient to induce long-term consequences (42). Future studies are necessary to determine if (+)-JQ1-mediated repression of LPS-induced increased chromatin accessibility at putative AP-1 and IRF motifs are permanent following removal of (+)- JQ1 from culture and should include re-challenge with LPS to assess these effects during macrophage tolerization (43).

In summary, our study provides evidence that BET inhibitors have the ability to overcome dysregulated inflammatory signaling in *Il10*^-/-^ macrophages and accomplishes this through prevention of bacterial stimulation dependent chromatin remodeling. Due to the heterogeneous nature of both CD pathogenesis and presentation, it will be necessary to isolate LP MΦs from CD patients to identify underlying chromatin aberrancies driving inflammation and determine if our results can be recapitulated in *ex vivo* co-culture systems prior to the pursuit of clinical trials.

## Materials and methods

### Mice

*Il10*^-/-^ mice on the C57BL/6J background maintained under specific pathogen free (SPF) conditions were used to generate BMDMs. Germ free *Il10*^-/-^ mice (C57BL/6J background) were bred and maintained by the National Gnotobiotic Rodent Resource Center (NGRRC) and used to evaluate the clinical efficacy of (+)-JQ1.

### Colonization and treatment of mice

8-12-week-old *Il10*^-/-^ mice (C57BL/6J background) were removed from germ free housing and placed in isolation in a BSL2 facility for the duration of these experiments. Mice were injected I.P. with 50 mg/kg (+)-JQ1 (MedChemExpress, #HY-13030) resuspended in 20% sulfobutylether-β-cyclodextrin (SBE-β-CD) (MedChemExpress, #HY-17031) or 10% DMSO diluted in 20% SBE-β-CD. Mice were subsequently colonized by oral and rectal swabbing with slurries made from feces of C57BL/6J mice raised under SPF conditions resuspended in pre-reduced anaerobic 1x PBS. Additional I.P. injections were given on Days 3, 6, 9, and 12. Weight was measured daily through to harvest on Day 14.

### Macrophage isolation and stimulation

BMDMs from 12-week-old mice were harvested and cultured as previously described in biological triplicate (one animal per replicate, aged 12 weeks) (9). BMDMs were counted and re-plated in duplicate for RNA- and ATAC-seq experiments. Macrophages were treated with 500nM (+)-JQ1 (MedChemExpress #HY-13030), 10 ng/mL recombinant IL-10 (PeproTech, #210-10), or vehicle control for 12 hours followed by the addition of 50 ng/mL LPS (InvivoGen, #tlrl-peklps) for 4 hours. Untreated, unstimulated *Il10*^-/-^ BMDMs served as an additional control and remained in culture for 16 hours. Samples were collected in TRIzol or freezing media for RNA- and ATAC-seq experiments, respectively.

### RNA-seq

RNA was isolated from murine BMDMs using the Norgen Biotek Corp. Total RNA Purification Kit (Cat. 17200) according to the manufacturer’s protocols. These kits use column-based DNase treatment to eliminate DNA contamination.

RNA-seq libraries were prepared using the Illumina KAPA Stranded RNA-seq Kit with RiboErase (HMR). Paired-end (50bp) sequencing was performed on the Illumina HiSeq 4000 platform (Gene Expression Omnibus [GEO] accession no. pending). Reads were extracted using TagDust v1.13 and aligned to the mm9 genome assemblies using STAR with default parameters (9, 44, 45). Reads were quantified using RSEM v1.2.31 with default parameters (9, 46). PCA was performed using the prcomp function in R on DESeq2 normalized variance stabilizing transformation (VST) transformed counts for the top 1,000 most variably expressed genes (47).

Pairwise differential gene expression between LPS stimulated and unstimulated samples and between LPS stimulated, (+)-JQ1 treated and unstimulated samples was determined using the DESeq2 Wald test (47). To focus on genes significantly affected by (+)-JQ1 treatment during LPS stimulation, gene expression for individual biological replicates for LPS stimulated and LPS + (+)- JQ1 conditions were normalized to the average expression across unstimulated *Il10*^-/-^ BMDMs and DEGs (base mean expression ≥ 10 and FDR < 0.05) were identified using the likelihood ratio test (LRT) in DESeq2 (47). Distinct gene expression classes (I – X) were identified based on varying combinations of levels of statistical significance and log_2_ fold-change of individual Wald tests (Supplementary Table 1). Z-scores were calculated based on the log_2_ fold-change of individual biological replicates for LPS stimulated and LPS stimulated + (+)-JQ1 treated samples normalized to the average of the unstimulated and untreated *Il10*^-/-^ BMDMs. These z-scores were used to generate heatmaps of the ten classes. Enriched KEGG pathways (FDR < 0.05) were identified using Enrichr (48).

### ATAC-seq

ATAC-seq was performed as previously described (49). To prepare nuclei, cells were pelleted and washed with cold PBS followed by lysis using cold lysis buffer (10mM Tris-HCl, 10mM NaCl, 3 mM MgCl2, and 0.1% NP-40). Pelleted nuclei underwent transposition using the Nextera DNA Library Prep Kit (Illumina #FC-121-1030). Samples were resuspended in the transposition reaction (12.5μL 2x TD buffer, 2 μL transposase, and 10.5 μL nuclease-free water) and incubated at 37C for 1 hour at 1000 RPM. Transposed DNA samples were purified using the Qiagen MinElute Kit (#28204) followed by amplification using 1x PCR master mix (NEB #M0541S) and 25 μM Ad1_noMX and Ad2.* indexing primer for 10-14 cycles. Libraries were purified and size selected by magnetic separation using Agencourt AMPure XP magnetic beads (Beckman Coulter #A63880). Paired-end (50bp) sequencing was performed on the Illumina HiSeq 4000 platform (GEO accession no. pending). Data was processed using PEPATAC v0.9.0 with default parameters (50). Uniquely mapped reads were aligned to the mm9 genome using bowtie2 (51). Peaks were called on individual samples using MACS2 (52, 53). ChIPSeeker was used to classify peaks as promoter or distal, where any peak that that did not fall in the −1000bp/+100bp from the TSS of mm9 reference genes was classified as distal (54). BEDTools was used to determine distal peaks +/-25kb from the TSS (50kb window) of each gene (55).

PCA was performed using the prcomp function in R on DESeq2 normalized, variance stabilizing transformation (VST) transformed counts for the top 10,000 most variably accessible peaks (47). Promoter and distal peaks were calculated and scaled with the computeMatrix tool from DeepTools v 3.5.0 and plotted using either plotHeatmap or plotProfile (56). Differentially accessible regions (DARs) across the union set of peaks was determined using DESeq2 (47). DARs with a |log_2_ fold-change| > 1 were identified using VennDiagram v1.6.20 (57). VST transformed counts were used for Spearman-ranked correlation analyses.

For each gene, DARs +/-25kb of the TSS were associated with the gene. Enrichments of DARs around genes in expression-defined categories (I – X) were calculated using a hypergeometric test. Known transcription factor motif analysis was performed with *findMotifsGenome* from HOMER using the middle 500bp window of DARs (FDR < 0.05). The *annotatePeaks* command was used to identify the location of enriched transcription factor motifs (FDR < 0.10) (58, 59). Motif matrices were clustered using Regulatory Sequence Analysis Tools (RSAT) using the Ncor metric for motif-to-motif similarity matrix with the lower thresholds set to 5 for width, 0.7 for Pearson correlation, and 0.4 for relative width-normalized Pearson correlation (60).

### Histology

Slides containing sections of proximal and distal colon were prepared for H & E staining. Two independent scorers blinded to the experimental and control groups performed histological analysis using an established scoring system for evaluating goblet cell loss and submucosal infiltrate in animal models (61). Histological sub-scores were added together to generate a composite histology score.

### Tissue RNA extraction and qPCR

Total RNA was isolated from cecal tissue taken from GF (Day 0) and colonized (Day 14) *Il10*^-/-^ mice using the Norgen Biotek Corp. Total RNA Purification Kit (Cat. 17200). cDNA was derived from 500 ng RNA by reverse transcriptase using the High Capacity cDNA Reverse Transcription Kit (Applied Biosystems, 4368814) according to the manufacturer’s specifications. Quantitative real-time PCR was performed using 10 ng cDNA with the PowerUp SYBR Green Master Mix (Applied Biosystems, A25741) and 10 μM of forward and reverse primers for *Il12β* (F: 5’ CGCAAGAAAGAAAAGATGAAGGAG 3’ R: 5’ TTGCATTGGACTTCGGTAGATG 3’) and *β-actin* (F: 5’ AGCCATGTACGTAGCCATCCAG 3’ R: 5’ TGGCGTGAGGGAGAGCATAG 3’). Reactions were performed in triplicate. Fold-change was calculated by determining ΔΔCt for all reactions followed by normalization to the average ΔΔCt value of GF animals.

### Flow Cytometry

*Il10*^-/-^ BMDMs were cultured for 16 hours with 500 nM (+)-JQ1. Cells were washed with cold 1x PBS followed by staining with LIVE/DEAD Fixable Blue Dead Cell Stain Kit (1:1000; Invitrogen L23105). Cell cycle changes in (+)-JQ1 treated *Il10*^-/-^ BMDMs were assessed using the BD Pharmingen BrdU Flow kit according to the manufacturer (559619). Briefly, *Il10^-/-^* BMDMs were co-cultured with BrdU and treated with (+)-JQ1 for 16 hours followed by fixation and permeabilization. Cells were treated with DNase followed by staining with FITC-conjugated anti-BrdU.

Bulk LPMCs were isolated from the colons of *Il10*^-/-^ mice by an enzymatic method followed by Percoll density gradient as previously described (9). Splenocytes were isolated as previously described (9). LPMCs and splenocytes were washed with cold 1x PBS and stained for viability followed by cell-surface marker staining for CD45 (1:100; Clone 30-F11, BioLegend), CD3ε (1:300; Clone 145-2C11, BioLegend), CD19 (1:200; Clone 6D5, BioLegend), CD11b (1:200; M1/70, BD Biosciences), CD11c (1:200; Clone N418, BioLegend), and F4/80 (1:200; Clone BM8, BioLegend) diluted in staining buffer (5% FBS/PBS). All samples were fixed with 4% PFA. Data was acquired with the FACSDIVA software using the BD LSR II and analyzed using FlowJo version 10.7.1.

### Statistics

Differential analyses of ATAC-seq and RNA-seq data were performed using DESeq2 with FDR adjusted *P* values used to measure statistical significance (47). Spearman rank correlations of VST transformed counts were calculated using RStudio Version 1.2.5033. Known motif enrichment was determined using HOMER, which generates *P* values by screening a library of reliable motifs against target and background sequences. GraphPad Prism 8 for Mac (Graphpad Software Inc.) was used for all other statistical analyses. Statistical significance comparing the absolute log_2_ fold-change values for LPS stimulated and LPS stimulated + (+)-JQ1 treated samples were determined by Wilcoxon match-paired signed rank tests. Statistical differences in relative gene expression for the LPS, LPS + (+)-JQ1 and LPS + IL-10 conditions were determined using one-way ANOVA followed by post hoc analyses using Tukey’s multiple comparisons test. Paired parametric t-tests were used to determine statistical significance for (+)- JQ1 and DMSO treated BMDMs for the evaluation of the number of cells in S-phase and percentage of viable cells. All other statistics were determined using a 2-tailed unpaired, nonparametric, student’s t test. For all tests, except for motif analysis, *P*_adj_ < 0.05, empirical *P* < 0.05, or *P* < 0.05 were considered statistically significant. For motif analysis, *P*_adj_ < 0.1 was considered statistically significant.

### Study Approval

All animal experiments were performed in accordance with protocols approved by the Institutional Animal Care and Use Committee of the University of North Carolina at Chapel Hill (19-108.0).

## Supporting information

Supplemental Figures/Tables with legends

## Author Contributions

M. Hoffner O’Connor, T.S. Furey, and S.Z. Sheikh conceptualized the idea for the study and wrote the manuscript. M. Hoffner O’Connor, R.B. Sartor, T.S. Furey, and S.Z. Sheikh designed the experiments. M. Hoffner O’Connor, A. Berglind, Z.J. Lynch, M. R. Schaner, E.C. Steinbach, J. Herzog and O.K. Trad performed the experiments. M. Hoffner O’Connor, T.S Furey, M.M. Kennedy, and B.P. Keith analyzed the sequencing data. W.R. Jeck and J.C. Arthur scored the histology. S.Z. Sheikh and T.S. Furey supervised and funded the project.

## Acknowledgements

We gratefully acknowledge the National Gnotobiotic Rodent Resource Center for providing all of the germ free mice used for this study (NIDDK P40OD010995, & NIDDK P30DK034987, and Crohn’s and Colitis Foundation), the technical support from the UNC High Throughput Sequencing Facility (NCI P30-CA016086 & P30-ES010126), the Center for Gastrointestinal Biology and Disease Histology Core for processing and staining all histological specimens (NIDDK P30DK034987), and use of the LSRII by the UNC Flow Cytometry Core Facility (NCI P30-CA016086).

We also thank members of the Sartor Lab, including Muyiwa Awoniyi, Bo Liu, and Akihiko Oka for generously providing SPF *Il10*^-/-^ mice used for this study and for technical support for the *in vivo* experiments performed for this study, the Gulati lab for providing material for GF mouse colonization, and Tamara Vital for providing technical support associated with the preliminary ATAC-seq analyses and supportive conversations throughout the progression of this project.

