## Supplemental Figures/Tables with legends for "BET protein inhibition regulates macrophage chromatin accessibility and microbiota-dependent colitis"

### Supplementary Figures

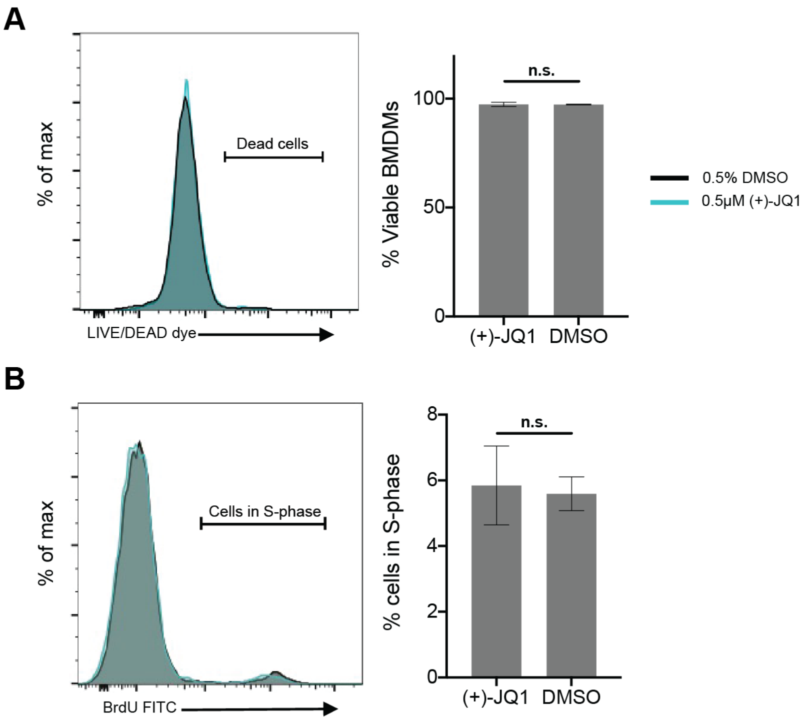


**Supplementary Figure 1. Pro-longed (+)-JQ1 treatment does not alter viability or cell cycle progression in BMDMs.** (A) Evaluation and quantification of (A) LIVE/DEAD dye or (B) BrdU FITC incorporation in BMDMs treated with (+)-JQ1 or 0.5% DMSO. N=2 biological replicates. Statistics generated using 2-tailed, paired, student's t-test.

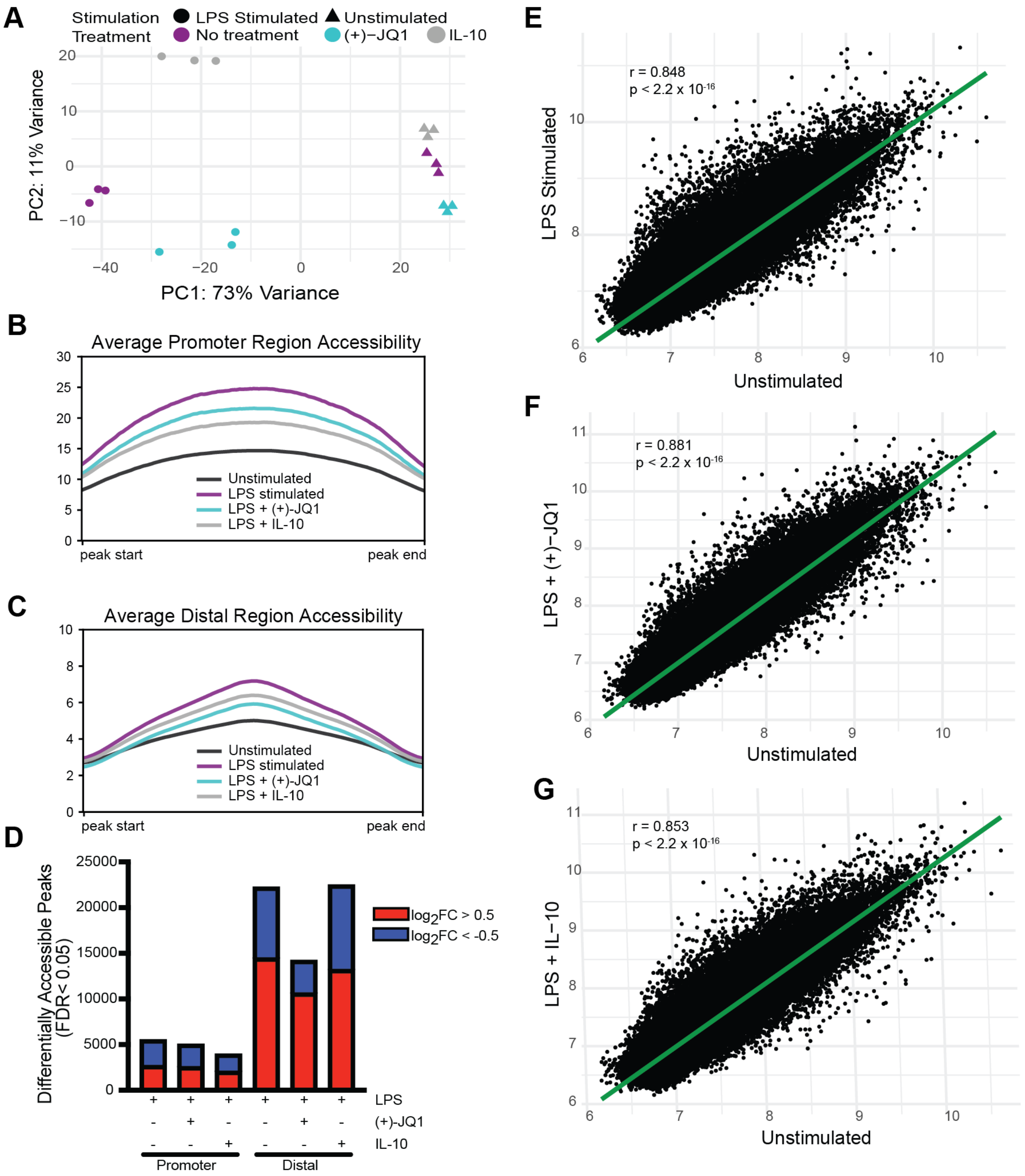


**Supplementary Figure 2. Chromatin remodeling with (+)-JQ1 treatment is distinct from exogenous IL-10.** (A) Principle component analysis (PCA) of chromatin accessibility profiles for samples that remained unstimulated (triangles) or were stimulated with LPS (circles) in the absence of treatment (purple) presence of (+)-JQ1 (aqua) or IL-10 (gray). Average chromatin accessibility signal profiles for (B) promoter regions (-1000/+100 bases) and (C) distal regions for unstimulated (black), LPS stimulated (purple), LPS + (+)-JQ1 (aqua) and LPS + IL-10 (gray) samples. (D) Identification of regions that differentially increase (red) or decrease (blue) in accessibility during LPS stimulation with or without (+)-JQ1 or IL-10 as compared with unstimulated *Il10*$^{-/-}$ BMDMs at promoter (left) and distal (right) regions (FDR < 0.05). Spearman rank correlations using VST transformed counts correlating unstimulated counts with (E) LPS stimulation (rho = 0.848), (F) LPS + (+)-JQ1 (rho = 0.881), and (G) LPS + IL-10 (rho = 0.853) conditions. FDR adjusted P values determined using DESeq2. Spearman rank correlations determined by RStudio. N=3 biological replicates.

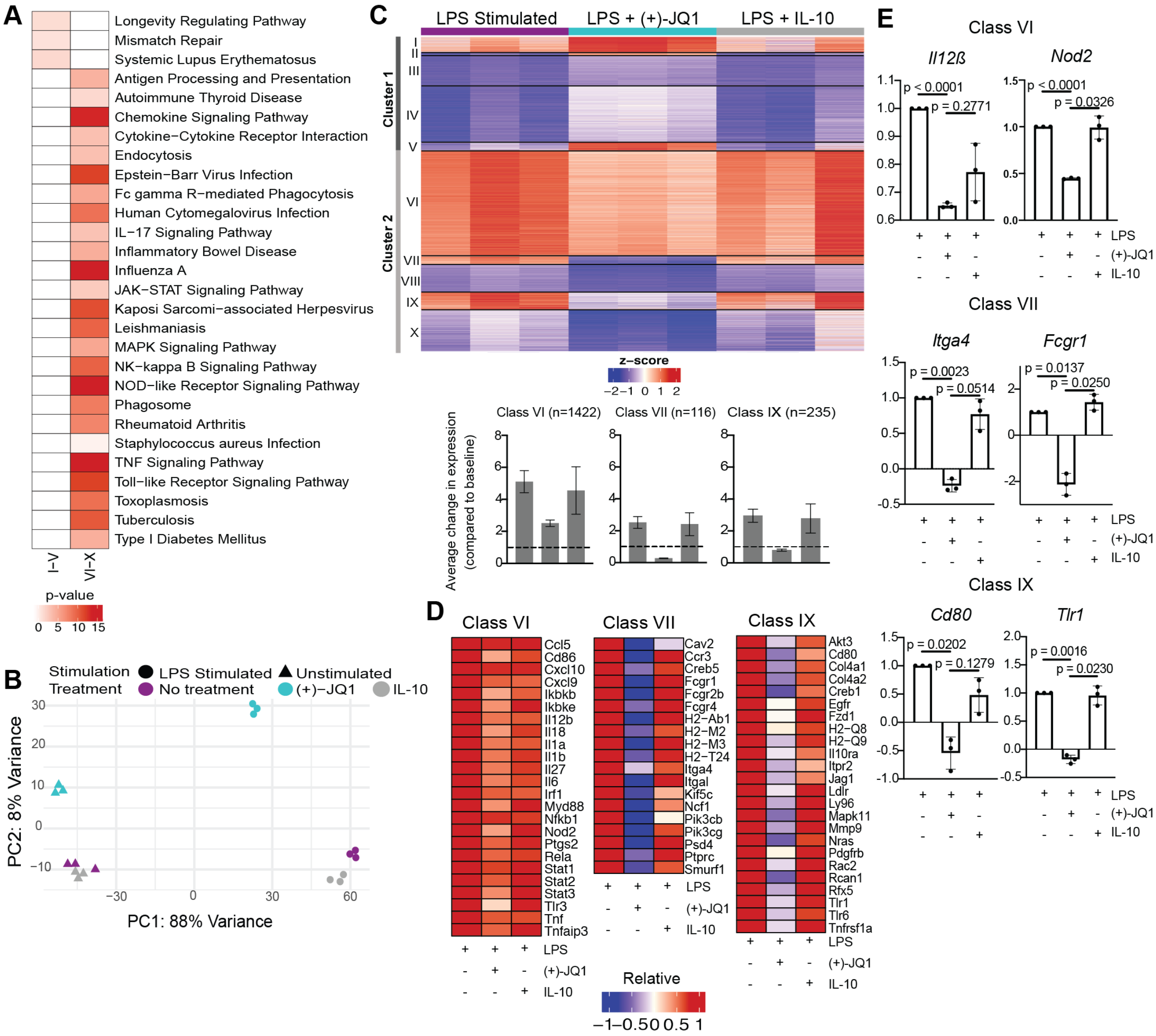


**Supplementary Figure 3. Exogenous IL-10 has limited impact on LPS-induced inflammatory gene expression as compared with (+)-JQ1.** (A) KEGG pathway analysis for all genes in cluster 1 (Classes I – V) and cluster 2 (Classes VI – X) (B) Principle component analysis (PCA) of gene expression profiles for samples that remained unstimulated (triangles) or were stimulated with LPS (circles) in the absence of treatment (purple) presence of (+)-JQ1 (aqua) or IL-10 (gray). (C) Representative heatmap of calculated z-scores based on $\log_2$ fold-change of individual replicates for the LPS, LPS + (+)-JQ1, and LPS + IL-10 conditions normalized to the average of unstimulated, untreated *Il10*$^{-/-}$ BMDMs. Genes were identified and divided into 2 major clusters and sub-divided into 5 classes through differential analyses outlined in Supplementary Table 1. Bar graphs represent average change in expression for LPS, LPS (+)-JQ1, and LPS + IL-10 conditions normalized to baseline for Classes. VI, VII, and IX. (D) Heatmaps of representative genes from Classes VI, VII, and IX that correspond to genes induced by LPS that are attenuated with (+)-JQ1 and compare to addition of IL-10. (E) Normalized expression of individual genes found in Classes VI, VII, and IX for the LPS stimulated, LPS + (+)-JQ1, and LPS + IL-10 conditions. Significance values determined using a one-way ANOVA followed by Tukey's multiple comparisons test. N=3 biological replicates.

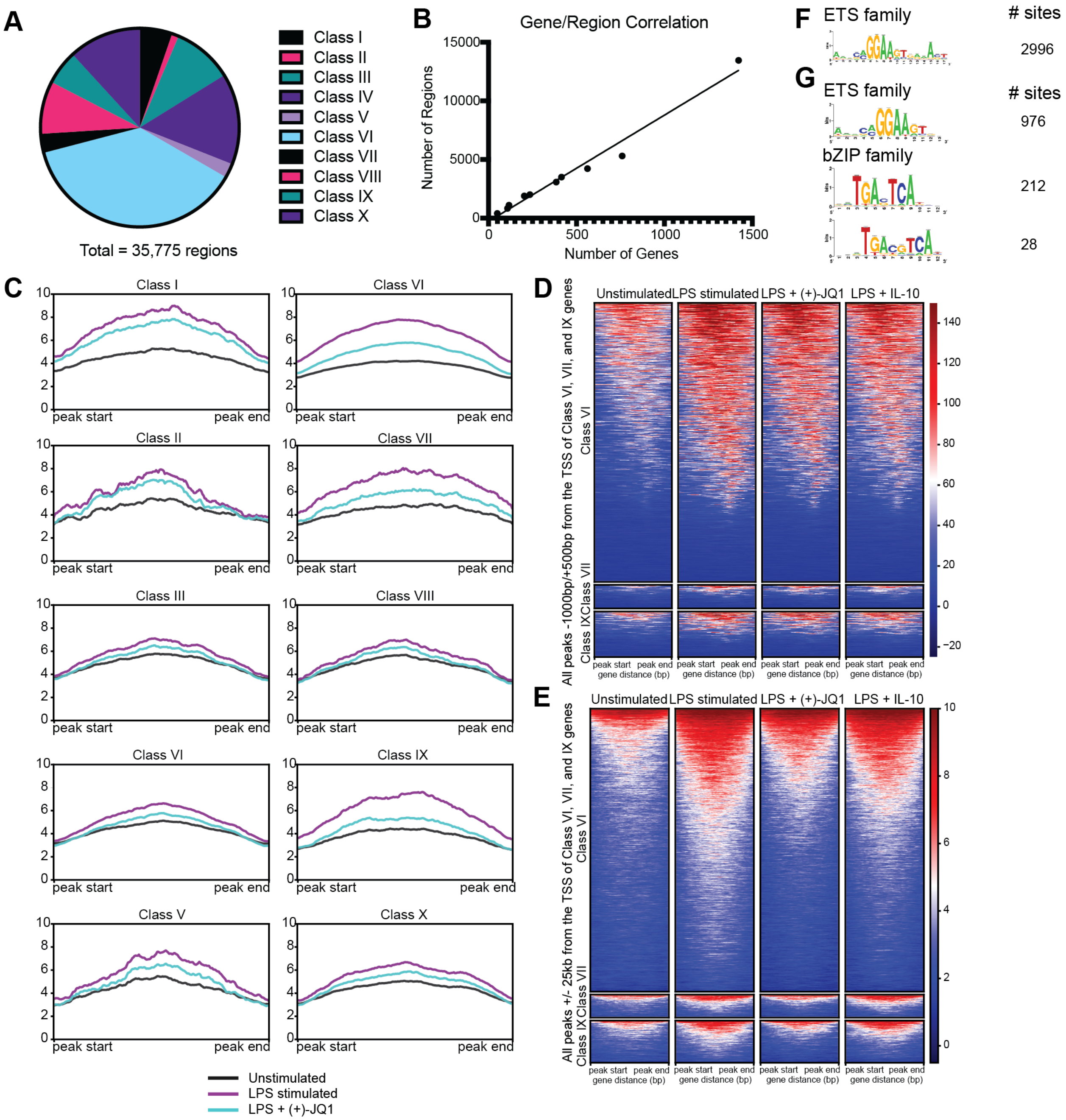


**Supplementary Figure 4. Lack of LPS-induced chromatin remodeling at distal regions associated with Class VI is unique to (+)-JQ1 treatment.** (A) Pie chart representing the number of distal peaks associated with each gene class using BEDtools intersect. (B) Correlation of the number of DEGs associated with each gene class and the number of peaks identified. (C) Average signal plots for all distal regions associated with Classes I – X. Heatmaps of accessible chromatin during unstimulated (left), LPS stimulated (center), LPS + (+)-JQ1, or LPS + IL-10 conditions for Classes VI (top), VII (center), and IX (bottom) for ChIPSeeker identified (D) promoter regions and (E) distal regions. Identification, motif clustering, and quantification of sites containing known HOMER motifs for regions that (F) decrease in accessibility with LPS stimulation as compared with unstimulated BMDMs or (G) increase with (+)-JQ1 treatment prior to LPS stimulation as compared with LPS stimulation alone. FDR < 0.05 for differential analysis;

FDR < 0.10 for motif enrichment. FDR adjusted P values for differential analysis determined using DESeq2 and for motifs using HOMER. N=3 biological replicates.

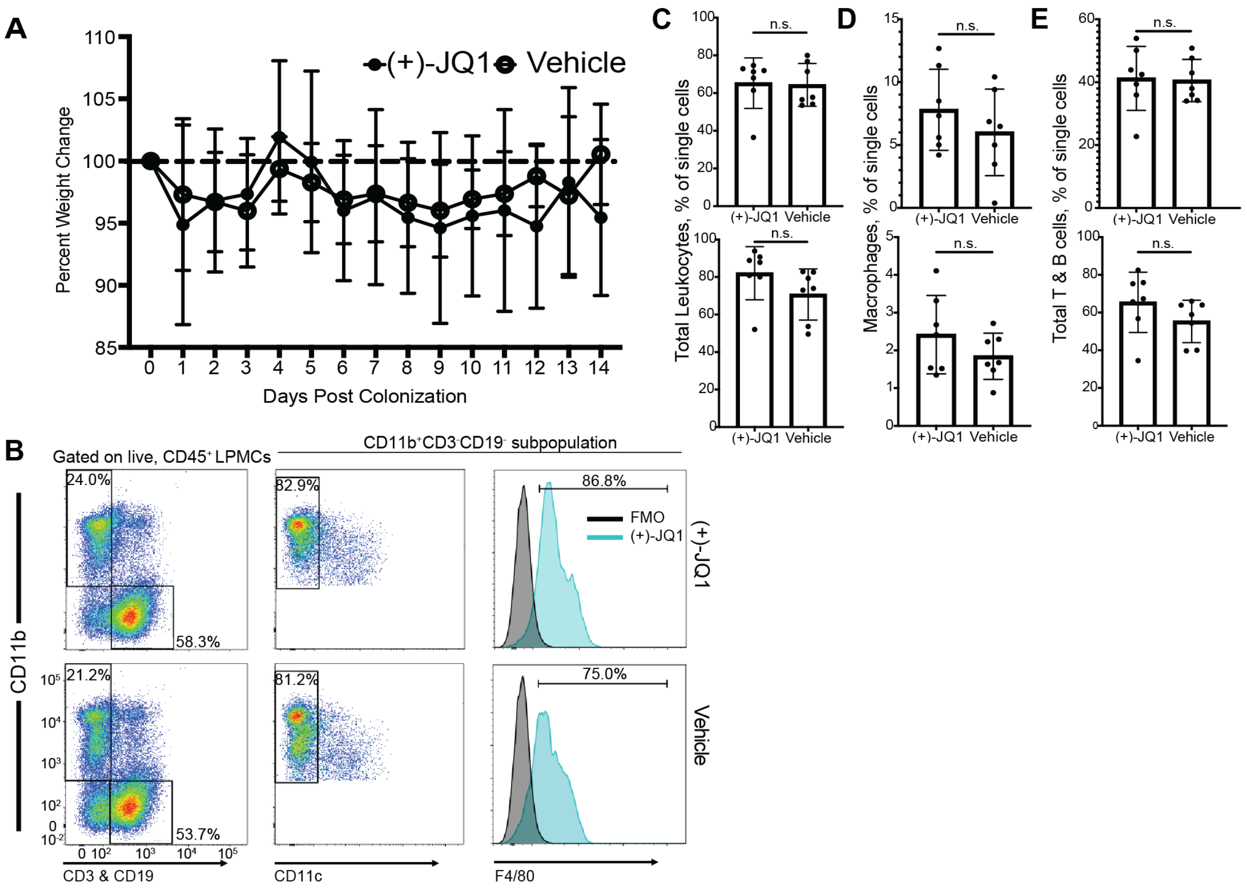


**Supplementary Figure 5. (+)-JQ1 treatment does not have a significant effect on bulk immune cell population size.** (A) Daily percent change associated with treatment using (+)-JQ1 or vehicle control (n = 16). (B) Representative gating strategy for $CD11b^{+}CD11c^{-}F4/80^{+}$ macrophage and $CD3^{+}/CD19^{+}$ lymphocyte populations bulk lamina propria mononuclear cells (LPMCs) isolated from murine colons. Quantification of colonic (top) and splenic (bottom) (C) total $CD45^{+}$ leukocytes (D) $CD11b^{+}CD11c^{-}F4/80^{+}$ macrophages and (E) $CD3^{+}$ & $CD19^{+}$ lymphocyte populations normalized to the number of single cells acquired by flow cytometry (n = 7/treatment). Statistics generated using two-tailed, unpaired, student's t test.

### Supplementary Tables

Supplementary Table 1. LRT and Wald (pair-wise) test parameters used to identify 10 gene expression classes with (+)-JQ1 treatment and LPS stimulation.

| Class | $Log_2FC$ Direction (LRT) | $P_{adj}$ (unstimulated vs LPS stimulated) | $Log_2FC$ (unstimulated vs LPS stimulated) | $P_{adj}$ (unstimulated vs LPS + (+)-JQ1) | $Log_2FC$ (unstimulated vs LPS + (+)-JQ1) | Number of Genes |
|---|---|---|---|---|---|---|
| I | $Log_2FC>0$ | $P_{adj}<0.05$ | $Log_2FC>0$ | $P_{adj}<0.05$ | $Log_2FC>1$ | 203 |
| II | $Log_2FC>0$ | $P_{adj}<0.05$ | $Log_2FC<0$ | $P_{adj}<0.05$ | $Log_2FC>1$ | 50 |
| III | $Log_2FC>0$ | $P_{adj}<0.05$ | $Log_2FC<0$ | $P_{adj}<0.05$ | $Log_2FC<-1$ | 415 |
| IV | $Log_2FC>0$ | $P_{adj}<0.05$ | $Log_2FC<0$ | $P_{adj}>0.05$ | NA | 760 |
| V | $Log_2FC>0$ | $P_{adj}>0.05$ | NA | $P_{adj}<0.05$ | $Log_2FC>1$ | 108 |
| VI | $Log_2FC<0$ | $P_{adj}<0.05$ | $Log_2FC>1$ | $P_{adj}<0.05$ | $Log_2FC>0$ | 1422 |
| VII | $Log_2FC<0$ | $P_{adj}<0.05$ | $Log_2FC>1$ | $P_{adj}<0.05$ | $Log_2FC<0$ | 116 |
| VIII | $Log_2FC<0$ | $P_{adj}<0.05$ | $Log_2FC<-1$ | $P_{adj}<0.05$ | $Log_2FC<0$ | 285 |
| IX | $Log_2FC<0$ | $P_{adj}<0.05$ | $Log_2FC>1$ | $P_{adj}>0.05$ | NA | 235 |
| X | $Log_2FC<0$ | $P_{adj}>0.05$ | NA | $P_{adj}<0.05$ | $Log_2FC<0$ | 563 |
